# Increased resistance of SARS-CoV-2 Omicron Variant to Neutralization by Vaccine-Elicited and Therapeutic Antibodies

**DOI:** 10.1101/2021.12.28.474369

**Authors:** Takuya Tada, Hao Zhou, Belinda M. Dcosta, Marie I. Samanovic, Vidya Chivukula, Ramin S. Herati, Stevan R. Hubbard, Mark J. Mulligan, Nathaniel R. Landau

## Abstract

Currently authorized vaccines for SARS-CoV-2 have been highly successful in preventing infection and lessening disease severity. The vaccines maintain effectiveness against SARS-CoV-2 Variants of Concern but the heavily mutated, highly transmissible Omicron variant poses an obstacle both to vaccine protection and monoclonal antibody therapies. Analysis of the neutralization of Omicron spike protein-pseudotyped lentiviruses showed a 26-fold relative resistance (compared to D614G) to neutralization by convalescent sera and 26-34-fold resistance to Pfizer BNT162b2 and Moderna vaccine-elicited antibodies following two immunizations. A booster immunization increased neutralizing titers against Omicron by 6-8-fold. Previous SARS-CoV-2 infection followed by vaccination resulted in the highest neutralizing titers against Omicron. Regeneron REGN10933 and REGN10987, and Lilly LY-CoV555 and LY-CoV016 monoclonal antibodies were ineffective against Omicron, while Sotrovimab was partially effective. The results highlight the benefit of a booster immunization in providing protection against Omicron but demonstrate the challenge to monoclonal antibody therapies.

## Introduction

The vaccines that have been granted emergency use authorization (EUA) have proven highly protective against SARS-CoV-2, resulting in a major decrease in infection rates, hospitalization and deaths [1]; however, the appearance of recently evolved viral variants classified as variants of concern (VOC) [2] that contain multiple mutations in the viral spike protein have raised concerns about potential decreases in vaccine effectiveness. These concerns have been assuaged by laboratory findings of modest 2-5-fold decreases in neutralizing antibody titer against the VOCs [2-7] and epidemiological evidence of continued vaccine protection [8, 9]. Vaccination has been found to provide 78% protection against infection by the Delta variant, 90% protection against hospitalization and 91% protection against death [10]. Vaccines with current EUA status include the BNT162b2 and Moderna mRNA-1273 mRNA-based vaccines and J&J Janssen Ad26.COV2.S adenovirus vector-vaccine. In addition to vaccination, monoclonal antibody therapies have proven effective in preventing hospitalization and death. Monoclonal antibody cocktails from Regeneron consisting of REGN10933 (Casirivamab) and REGN10987 (Imdevimab), and from Eli Lilly consisting of LY-CoV016 (Etesevimab) and LY-CoV555 (Bamlanivimab) have proven effective at decreasing the frequency of hospitalization of COVID-19 patients [11-13]. The GlaxoSmithKline/Vir Biotechnology monoclonal antibody VIR-7183 has been shown to decrease hospitalization and risk of death by 79% in adults at high risk and has been granted EUA authorization by the U.S. Food and Drug Agency for the treatment of COVID-19 [15].

The identification of the newly emergent Omicron (B.1.1.529) SARS-CoV-2 variant has raised concerns about possible reductions in vaccine effectiveness. The variant was identified in COVID-19 patients in Botswana in early November, 2021 where it rapidly rose to a prevalence of 71% and was shortly thereafter identified in infected individuals in South Africa[16]. Prevalence of the Omicron variant has continued to increase rapidly as a result of the increased transmissibility of the virus, having now replaced Delta as the predominant variant in the U.S. with a current prevalence of 73.2% and up to 90% in metropolitan areas. While the vaccines have proven effective against earlier VOCs, the large number of mutations in the Omicron spike protein present the possibility of decreased antibody neutralizing titers against the Omicron variant, which could result in decreased protection from infection and disease.

As compared to the previously designated VOC spike proteins that contain 9-11 missense mutations, the Omicron spike protein has 34, 20 of which have not been found in previous VOCs or variant of interests (VOIs). These include 15 mutations in the receptor binding domain (RBD), 8 of which lie in the receptor binding motif (RBM) that directly contacts the receptor. The amino-terminal domain (NTD) has 8 mutations, 3 of which are deletions and one is a 3 amino acid insertion. The carboxy-terminal CTD has 10 mutations, 4 of which are close to the furin proteolytic processing site and three of which are close to the secondary processing site. The concomitant appearance of the multiple mutations in the Omicron virus suggests that some may have arisen from recombination with a related β-coronavirus or from extended replication in a chronically infected immunodeficient individual [17].

The large number of mutations in the Omicron RBD and NTD, which are the primary sites targeted by neutralizing antibodies, raises the possibility that the variant may be resistant to neutralization by current EUA approved vaccine-elicited antibodies, resulting in decreased protection from infection. It also raises the possibility that individuals previously infected with an earlier version of the virus might not be protected against re-infection by the Omicron variant. In addition, it raises a concern that the heavily mutated Omicron RBD might cause the failure of therapeutic monoclonal antibodies currently in clinical use to neutralize the virus, decreasing the effectiveness of their use in the treatment of severe COVID-19.

In this study, we used spike protein-pseudotyped lentiviral particles to measure the sensitivity of the Omicron variant to neutralization by vaccine-elicited antibodies in the sera of both naïve and recovered individuals and analyzed the neutralizing activity of the widely used therapeutic monoclonal antibodies. We found that the Omicron spike protein-pseudotyped virus was highly resistant to neutralization by the serum antibodies of individuals fully vaccinated with two immunizations of the Pfizer or Moderna mRNA vaccines. A homologous booster vaccination for individuals fully vaccinated with an mRNA vaccine increased neutralizing antibody titers 5-fold to a level predicted to provide a high degree of protection. Of concern, the monoclonal antibodies that constitute the Regeneron and Eli Lilly cocktails failed to neutralize virus with the Omicron spike protein. Sotrovimab was partially effective against the Omicron-pseudotyped virus.

## Material and Methods

### Plasmids

Plasmid expression vectors used in the production of lentiviral pseudotypes pMDL, pcVSV.G, pRSV.Rev and the lentiviral dual reporter virus genome pLenti.GFP.nLuc have been previously described [17]. The SARS-CoV-2 Omicron spike expression vector pc.Δ19.Omicron was chemically synthesized in two fragments encoding the codon-optimized open reading frame overlapping by 50 bp. The full-length coding sequence was generated by overlap extension PCR with the two fragments amplified with external primers containing a Kpn-I and Xho-I sites and then cloned into pcDNA6. Expression vectors encoding spike proteins with the individual mutations of the Omicron spike protein were generated by overlap extension PCR mutagenesis using the D614G spike protein plasmid pcCOV2.Δ19.D614G as template.

### Cells

293T, ACE2.293T and Vero cells were grown in Dulbecco’s Modified Eagle’s Medium/10% fetal bovine serum at 37°C under 5% CO_2_.

### Human sera and monoclonal antibodies

Human sera were collected at the NYU Vaccine Center with written consent of participants under IRB-approved protocols 18-02035 and 18-02037. Sera from convalescent were collected 32-57 days post-symptom onset. Sera from Pfizer BNT162b2-vaccinated, Moderna mRNA-1273-vaccinated study participants which were shown in Figure 1B were collected 90 and 80 days mean post-second immunization, respectively. Serum samples from study participants previously infected and subsequently vaccinated with BNT162b2 mRNA vaccine which shown in Figure 1C and 1D were collected 1 month and 7-8 months days post-second immunization. Sera from study participants vaccinated with BNT162b2 mRNA third boost vaccine were collected 1-month post-vaccination. Previous infection was documented by COVID-19 symptoms and a positive PCR test or serology.

**Figure 1.**
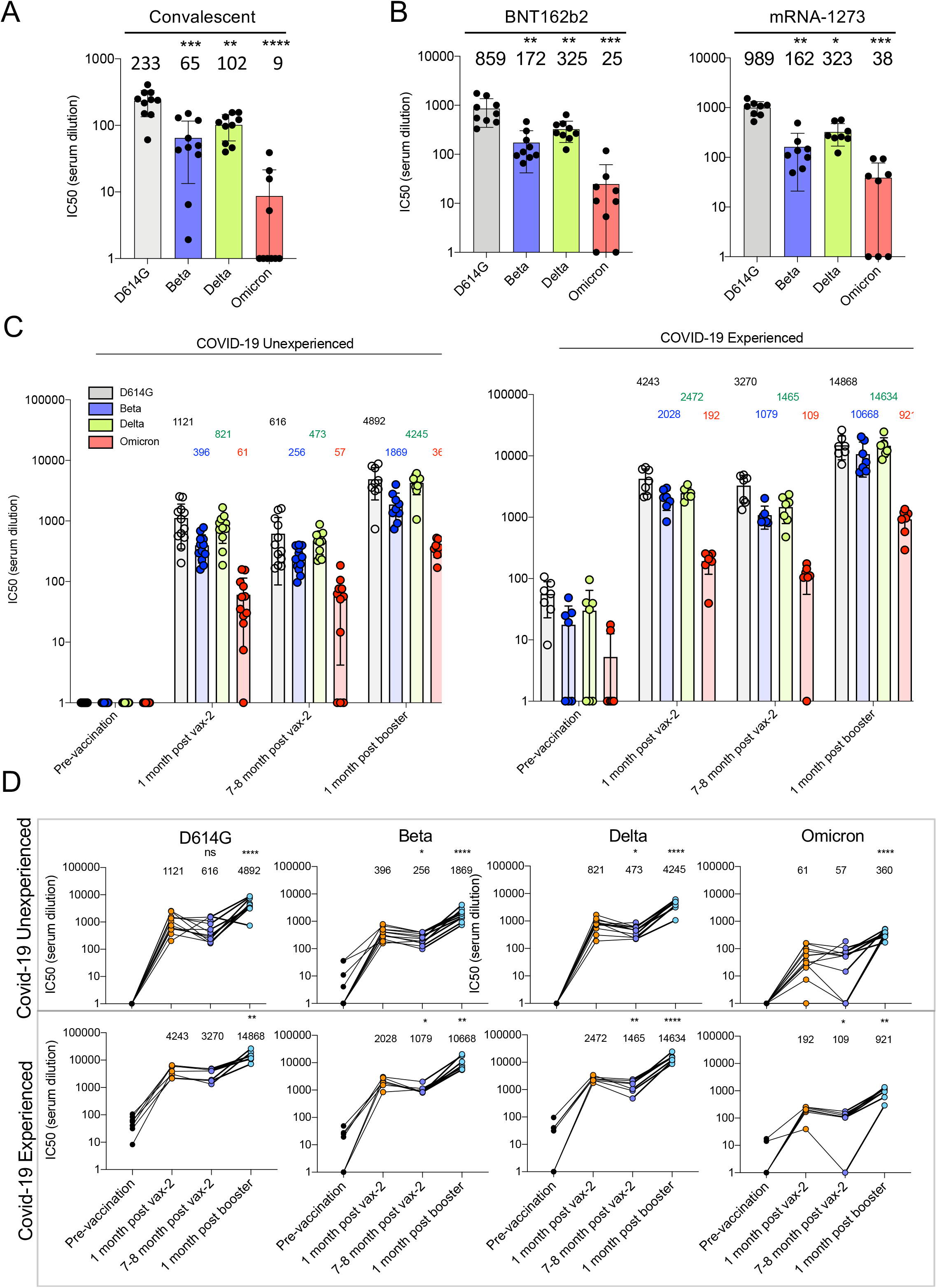
Decreased neutralization of Omicron spike protein-pseudotyped viruses by convalescent sera, mRNA vaccine-elicited antibodies. D614G, Beta, Delta and Omicron spike protein-pseudotyped viruses expressing dual GFP/nanoluciferase reporter genes with codon-optimized spike proteins deleted for the carboxy-terminal 19 amino acids were prepared as previously described [18]. Equivalent amounts of virus were mixed with a 2-fold serial dilution of donor serum and then applied to ACE2.293T cells. Luciferase activity was measured two days post-infection. Each serum dilution was measured in triplicate and the experiment was done twice with similar results and IC50 was determined. Statistical significance was calculated by two-sided testing. (*P≤0.05, **P≤0.01, ***P≤0.001, ****P≤0.0001). A. Neutralizing antibody titers of sera from study participants who had recovered from infection prior to the appearance of the current VOCs was measured against viruses pseudotyped by current VOCs (n=10). IC50 of each donor serum is shown with the Geometric mean titer (GMT) for each group shown above the bar. B. Neutralization of variant spike protein pseudotyped viruses by the sera of study participants fully vaccinated (two immunizations) with Pfizer BNT162b2 (n=9) and Moderna mRNA-1273 mRNA vaccines (n=8). C. Neutralizing antibody titers of study participants without or with a previous history of SARS-CoV-2 infection were measured on the pseudotyped viruses. Sera were collected from study participants pre-vaccination, 1-month post-second vaccination with Pfizer BNT162b2, 7-8 months post-second vaccination, and 1-month post-boost. Study participants were without previous SARS-CoV-2 infection (left) (n=12) or previously infected (right) (n=7). COVID-19 history was determined by symptoms and a PCR+ test or serology. GMTs for each group are shown above the bar. D. Sequential neutralizing antibody titers of sera from individual study participants without or with previous history of SARS-CoV-2 infection is shown for each of the study participants shown above in C. GMTs are shown above.

### SARS-CoV-2 spike protein lentiviral pseudotypes

Spike protein pseudotyped lentiviruses were produced by cotransfection of 293T cells with pMDL Gag/Pol packaging vector, lentiviral vector plenti.GFP.nLuc and spike protein expression vectors encoding 19 amino acid cytoplasmic tail deletions, as previously reported [17]. Transfected cell supernatants were harvested two days post-transfection and concentrated by ultracentrifugation. The viruses were normalized for reverse transcriptase (RT) activity and frozen in aliquots at -80°C.

### Antibody neutralization assay

Sera or monoclonal antibody was serially two-fold diluted and then incubated with an amount of virus corresponding to a volume that resulted in MOI=0.2 on ACE2.293T or Vero cells for pseudotyped virus. After 30-minute incubation at room temperature, the virus was added to 1 × 10^4^ target cells in a 96 well culture dish. The cells were cultured for 2 days after which the culture medium was removed and 50μl Nano-Glo luciferase substrate (Nanolight) was added. Luminescence was read in an Envision 2103 microplate luminometer.

### Data analysis

All samples were tested in duplicate or triplicate. Data were analyzed using GraphPad Prism 8 software and statistical significance was determined by the two-tailed unpaired t-test or nonparametric ANOVA test. Significance was based on two-sided testing and attributed to p< 0.05. Confidence intervals are shown as the mean ± SD or SEM (*P≤0.05, **P≤ 0.01, ***P≤ 0.001, ****P≤0.0001). Analyses of the structures of the SARS-CoV-2 spike protein with antibody Fabs was performed with the PyMOL Molecular Graphics System, v2.1.1 (Schrödinger, LLC).

## Results

### Increased resistance of virus with the Omicron spike to serum antibodies elicited by natural infection and vaccination

To determine the effectiveness of antibodies induced by infection with earlier SARS-CoV-2 variants to protect from re-infection with the Omicron variant, we tested neutralizing antibody titers in the sera of unvaccinated participants involved in an ongoing clinical study that had been collected 32 to 57 days post-COVID-19 symptom onset. Neutralizing antibody titers were measured using lentiviral virions pseudotyped by the parental D614G, Alpha, Beta and Delta spike proteins, an assay that accurately reflects titers obtained in the plaque reduction neutralization test (PRNT). The results showed modest reductions in neutralizing titer against Beta and Delta as compared to the parental D614G but a more substantial average 26-fold reduction in titer against Omicron. Approximately 60% of the donor sera had titers below the IC50 of 20 limit of detection in the assay **(Fig. 1A)**. To determine the effectiveness of antibodies elicited by vaccination, we tested sera collected 70 days post-immunization from study participants who had been fully vaccinated (two immunizations) with BNT162b2 or Moderna mRNA-1273 mRNA vaccines **(Fig. 1B)**. Titers against the D614G virus were 3-4-fold higher than those of the convalescent patient sera and the general pattern of neutralization of the variants was similar. Notably, neutralizing antibody titers against the Omicron pseudotype was decreased 26-34-fold compared to D614G.

Previous infection has been shown to strengthen and broaden the neutralizing antibody response to SARS-CoV-2 variants upon vaccination. To determine whether previous infection would increase neutralizing antibody titers against the Omicron variant, we tested sera from study participants who were vaccinated with BNT162b2 and had, or had not, been previously infected with SARS-CoV-2 **(Fig. 1C)**. Sera from study participants without previous infection, collected one month post-second vaccination, had high titers of neutralizing antibody against D614G virus; titers against Beta compared to D614G were decreased 2.8-fold, against Delta 1.4-fold and against Omicron 18-fold. Titers were had only slightly declined 7-8 months post-vaccination. One-month post-boost, titers increased for all variants. Titers against Omicron remained 14-fold lower than against D614G. Notably, study participants who had poor neutralizing titers against Omicron after two immunizations had increased their titers following the boost **(Fig. 1D)**. Sera from previously infected study participants were on average 3-4-fold higher than those without previous infection and had a similar ratio of neutralizing titers among the different variants. Sera from previously infected study participants post-boost achieved high neutralizing titers against the Beta and Delta variants. While titers against Omicron also rose, they remained 16-fold lower on average than that of D614G virus (14,868 for D614G; 921 for Omicron).

### Virus with the Omicron spike protein is resistant to therapeutic monoclonal antibodies

The Regeneron monoclonal antibody cocktail used for the treatment of COVID-19 consists of REGN10933 (Casirivamab) and REGN10987 (Imdevimab); the Eli Lilly and Company cocktail consists of LY-CoV016 (Etesevimab) and LY-CoV555 (Bamlanivimab). In addition, VIR-7831 (Sotrovimab) from GlaxoSmithKline and VIR Biotechnology has recently been given EUA approval. To determine the sensitivity of the Omicron variant to the therapeutic monoclonal antibodies, we analyzed their neutralizing titers against the D614G, Beta and Omicron spike protein pseudotyped viruses. REGN10933 potently neutralized D614G and Delta, was less active against Beta but had no detectable activity against Omicron **(Fig. 2A)**. REGN10987 also potently neutralized the earlier viruses but lacked activity against Omicron virus as did the REGN10933/REGN10987 cocktail. LY-CoV555 neutralized D614G and Alpha virus, had weak activity against Beta and Delta but was inactive against the Omicron virus **(Fig. 2B)**. LY-CoV016 potently neutralized the earlier viruses but lacked activity against Omicron virus as did the combined LY-CoV555/LY-CoV016 cocktail. VIR-7831 was active against Omicron but its IC50 was around 172-fold lower than against D614G **(Figure 2C)** and lower still when compared to the IC50 of the other monoclonal antibodies against the D614G virus. IC50s calculated from the curves in Figures 2A and B are shown in **Figure 2D**.

**Figure 2.**
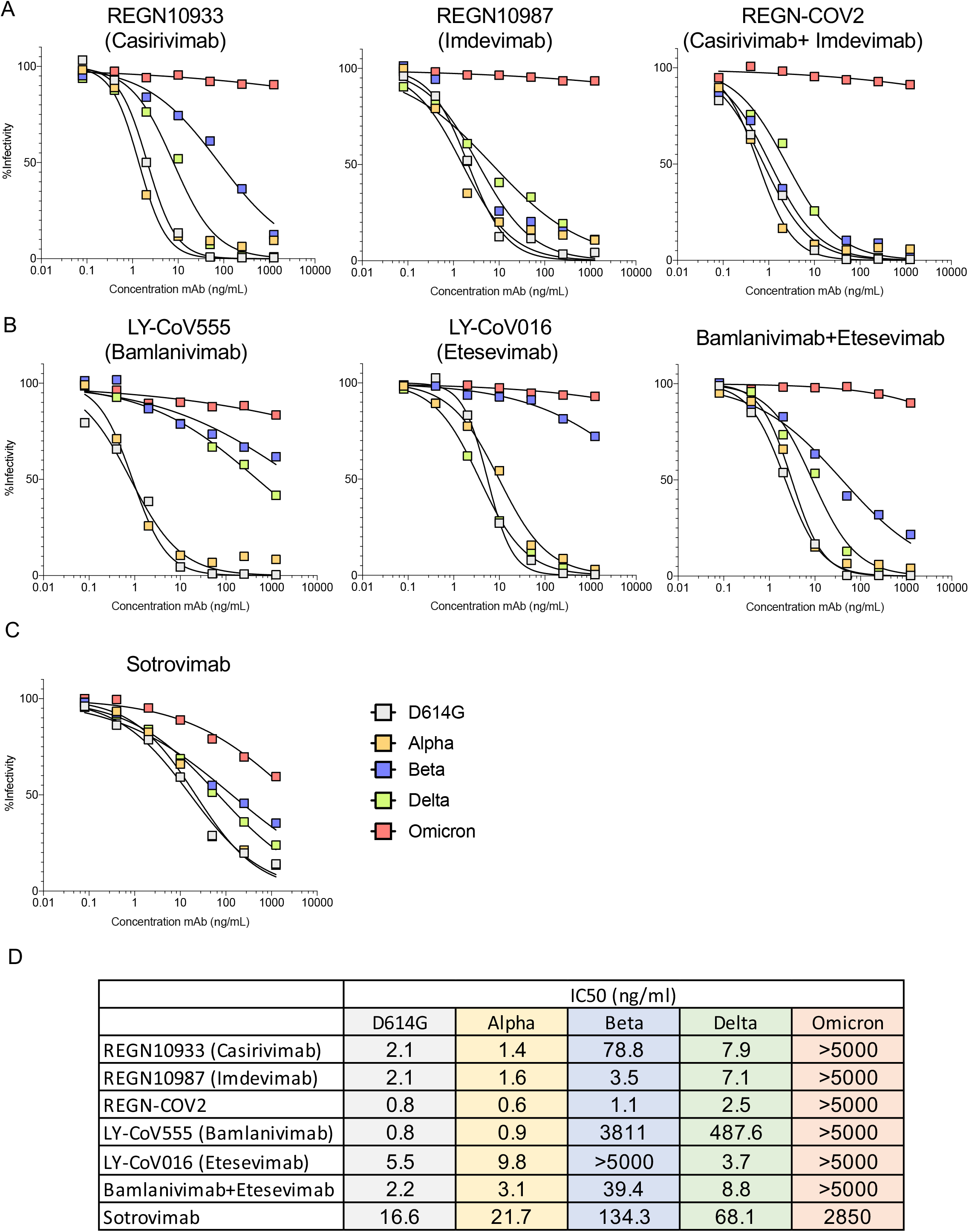
Therapeutic monoclonal antibodies have lost neutralizing activity against virus with the Omicron spike protein. A. Neutralization of viruses with the VOC spike proteins by Regeneron REGN10933 and REGN10987 monoclonal antibodies and the REGN-CoV-2 cocktail was measured using spike variant spike protein-pseudotyped viruses. B. Neutralization of viruses pseudotyped by the VOC spike proteins by LY-CoV555 (Bamlanivimab) and LY-CoV016 (Etesevimab) monoclonal antibodies was measured as in A above. C. Neutralization of viruses pseudotyped by the VOC spike proteins by VIR-7831 (Sotrovimab) was measured as in A above. D. The table shows the IC50s of the therapeutic monoclonal antibodies calculated using the data from the antibody neutralization curves shown in A, B and C. Larger numbers indicate decreased neutralization potency.

To determine which of the Omicron spike protein mutations allowed escape from neutralization, we tested the therapeutic monoclonal antibodies against a panel of viruses pseudotyped by spike proteins with the individual mutations of the Omicron RBD **(Fig. 3A)**. While most of the single mutations had no effect, specific mutations significantly decreased monoclonal antibody inhibitory activity (increased IC50). REGN10933 activity was affected by mutations K417N, E484A and Q493K **(Fig. 3B and 3C)**. REGN10987 was affected by mutations S371L, S373P, N440K, G446S with minor effects of several other mutations. The REGN10933/REGN10987 cocktail maintained most of its neutralization potency against the single point mutated virus. Etesevimab was affected by K417N, Q493K, Q498R and N501Y. Bamlanivimab inhibitory activity was ablated by mutations E484A and Q493K, while several other mutations had small effects. With the exception of E484A, most of the mutations had modest effects on neutralizing titer suggesting that the loss of activity by the monoclonal antibodies results from the combined effect of the full complement of Omicron spike protein mutations.

**Figure 3.**
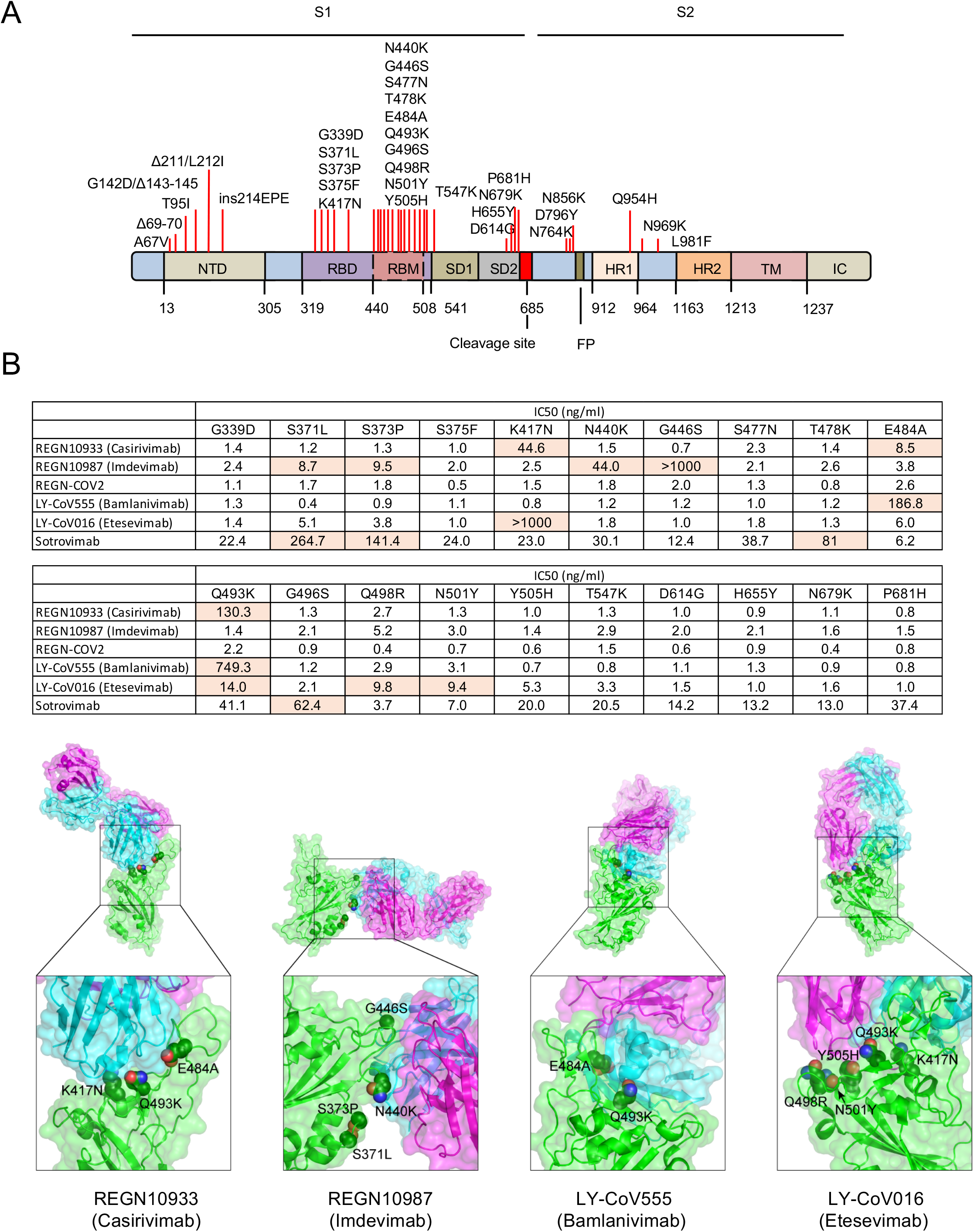
Omicron spike protein mutations that cause escape from therapeutic monoclonal antibodies are located at the antibody interaction interface. A. The location of Omicron mutations on the spike protein is diagrammed. The location of the S1 and S2 subunits of the processed spike protein, NTD, RBD, SD1, SD2, HR1, HR2, TM and IC domains are shown. Amino acid positions of the domains are labeled below. The furin cleavage site and hydrophobic fusion peptide (FP) are indicated. B. The table shows the IC50s calculated from the neutralization curves shown in Supplementary Figure 1. Mutations that caused >5-fold increase in IC50 are highlighted. C. The structures of Fabs from neutralizing antibodies bound to the SARS-CoV-2 spike protein are shown. For each antibody, the spike protein monomer is colored green, the Fab light chain is magenta, and Fab heavy chain is cyan. Mutations that adversely affect the neutralizing efficacy of each antibody are labeled, with the side chains of the D614G spike protein amino acid residues shown in sphere representation. Carbon atoms are colored green, oxygen atoms red, and nitrogen atoms blue. The PDB accession codes for the structures are 6XDG (Casirivimab and Imdevimab), 7KMG (Bamlanivimab), and 7C01 (Etesevimab).

The published crystal and cryo-electron microscopy structures of Fabs from neutralizing antibodies bound to the SARS-CoV-2 spike protein provide insights into how mutations in the Omicron spike protein interfere with antibody binding **(Figure 3C)**. The efficacy of Casirivimab (REGN10933) is compromised by mutations K417N, E484A, and Q493K. In the structure of the Casirivimab Fab-spike protein, these mutations are situated in the interface with the Fab heavy chain (Figure 3C). K417N would cause a loss of hydrogen bonding with T28 and T102 of the heavy chain, as well as the loss of a favorable electrostatic interaction with D31 of the heavy chain. E484A would result in the loss of hydrogen bonding with Y53 and S56 of the heavy chain, and Q493K would result in the loss of hydrogen bonding with N74 of the heavy chain.

For Imdevimab (REGN10987), four mutations in the Omicron spike protein lead to significant loss of efficacy: S371L, S373P, N440K, and G446S. The N440K mutation would create steric clashes between K440 and the heavy and light chains and result in charge repulsion with K55 of the light chain. Mutation of G446 to any other (larger) residue (e.g., G446S) would cause a steric clash with N57 of the heavy chain. Mutation of amino acids S371 and S373 adversely affect antibody activity but do not directly contact the Fab (Figure 3C); mutation of these amino acids could alter the stability of this loop segment, affecting the conformation of the nearby region (N440) of antibody binding.

For Bamlanivimab (LY-CoV555), E484A would result in the loss of salt bridges with R50 of the heavy chain and R96 of the light chain. Q493K would result in loss of a hydrogen bond with R104 of the heavy chain, and, critically, a lysine at this position would cause a steric and electrostatic clash with R104 of the heavy chain.

For Etesevimab (LY-CoV016), K417N would result in the loss of a salt bridge with D104 of the light chain, Q493K would result in the loss of a hydrogen bond with Y102 of the heavy chain, and lysine at this position would cause a steric clash with Y102. An arginine at position 498 (Q498R) would cause charge repulsion with R31 of the light chain. N501Y would be predicted to destabilize the local conformation of the spike protein, and tyrosine at this position would cause a steric clash with S28 of the light chain.

## Discussion

The emergence of the Omicron variant represents an evolutionary leap by SARS-CoV-2 in which 15 mutations were introduced into the RBD along with mutations and deletions in the NTD and CTD. As a result, the Omicron variant has developed resistance to neutralization by the serum antibodies of recovered individuals who had been infected with earlier SARS-CoV-2 variants to a degree that is expected to increase the number of individuals who become re-infected. In addition, virus with the Omicron spike has a high degree of resistance to neutralization by vaccine-elicited antibodies. The resistance might be expected given that current EUA approved vaccines encode the earlier D614G spike protein. While Alpha, Beta, Gamma and Delta VOCs show about a 3-4-fold resistance to neutralization by vaccine-elicited antibodies [2-7], virus with the Omicron spike protein has increased its resistance to neutralization by the serum antibodies of individuals fully vaccinated with BNT162b2 or Moderna-1273 by about 2640-fold, resulting in titers that are predicted by mathematical modeling to cause an increased frequency of breakthrough infections [19, 20].

Homologous boosting of SARS-CoV-2-inexperienced individuals by immunization with the Pfizer BNT162b2 vaccine increased neutralizing antibody titers against Omicron to levels that are predicted to be highly protective, although the titers remained about 10-fold below those against the other VOCs post-boost and the durability of the titers remains to be determined. Booster immunization of SARS-CoV-2 experienced individuals resulted in neutralizing antibody titers against Omicron approaching an IC50 of 1000, which as predicted by modeling will provide 90% protection against infection.

Our findings on monoclonal antibody neutralization of the Omicron variant suggest that the monoclonal antibodies currently in widespread use may become ineffective. REGN10933 (Casirivimab) and REGN10987 (Imdevimab) that constitute the Regeneron cocktail [18, 19] and LY-CoV555 (Bamlanivimab) [20, 21] and LY-CoV016 (Etesevimab) [22, 23] that constitute the Eli Lilly cocktail all failed to neutralize the virus. The recently approved VIR-7831 (Sotrovimab) [24] had significant neutralizing activity against virus with the Omicron spike protein although this was significantly decreased compared to titers against the other VOCs. Sotrovimab was 172-fold less active against the Omicron virus compared to the D614G virus. While neutralizing activity is considerably decreased, in treated patients, the antibody achieves a concentration of 24 μg/ml following a 500 mg dose. This concentration is well above the IC50 of Sotrovimab determined in tissue culture, and thus the antibody may prove beneficial for the treatment of COVID-19.

Mapping of the amino acid residues responsible for the escape from the monoclonal antibodies showed that most of the mutations had no effect but that several had partial effects on neutralization. The only mutation that had a dramatic effect was E484A, which ablated neutralization by LY-CoV555. The other mutations that compromised antiviral activity had modest effects. Thus, it was the cumulative effect of several mutations that abrogated antiviral activity for the other monoclonal antibodies. REGN10933, the neutralizing activity of which has been previously found to be affected by E484K and K417N of the Beta spike protein [3, 25, 26], is decreased another 8-fold by E484A of Omicron. REGN10987, which is nearly impervious to mutations in the earlier VOCs, was compromised by the constellation of five of Omicron mutations (S371L, S373P, N440K, G446S). K417N had a major effect (40-fold) on the activity of Etesevimab. The findings suggest that the Regeneron and Eli Lilly cocktails will not be effective for the treatment of patients infected by the Omicron variant. The effectiveness of Sotrovimab cannot be predicted from these data but it would seem likely that it will not be as effective on patients infected with Omicron variant as compared to those with Delta or other variants.

Our findings suggest that while the frequency of infections with the Omicron variant are likely to increase, the titers achieved by full vaccination followed by a booster immunization will protect most individuals from developing severe disease. The T cell response induced by vaccination, which is less prone to immune escape, may also provide additional protection. Our findings provide further support for the benefits of booster immunization and point to the need to develop additional therapeutics for the treatment of COVID-19.

The emergence of the Omicron variant raises concern about the possibility of additional evolutionary leaps for the virus and the need to preempt any such variants before they emerge. While the current surge in Omicron infections may increase hospitalization and mortality, there is also an increased likelihood of leading to herd immunity that may protect against future variants. The inclusion of additional antigens in the vaccines to further increase the T cell response may also prove beneficial in this regard.

### Study Limitations

This study was done on a relatively small number of participants which limits the resolution of fine difference in antibody titers in the different groups. In addition, it depends entirely on pseudotyped viruses rather than antibody neutralization of live virus. While pseudotyped virus has been shown to provide similar data to that of the live virus assays, it is conceivable that there could be differences [30].

## Acknowledgements

The work was funded by grants from the NIH to N.R.L. (DA046100, AI122390 and AI120898) and to M.J.M. (UM1AI148574).

## Author contributions

T.T. and N.R.L. designed the experiments. T.T., H.Z., B.M.D. and V.C. carried out the experiments and analyzed data. S.R.H. provided protein structural analyses. T.T., H.Z. and N.R.L. wrote the manuscript. M.I.S., R.H. and M.J.M supervised specimen selection and the collection of clinical information.

## Figure Legends

**Supplementary Figure 1.**
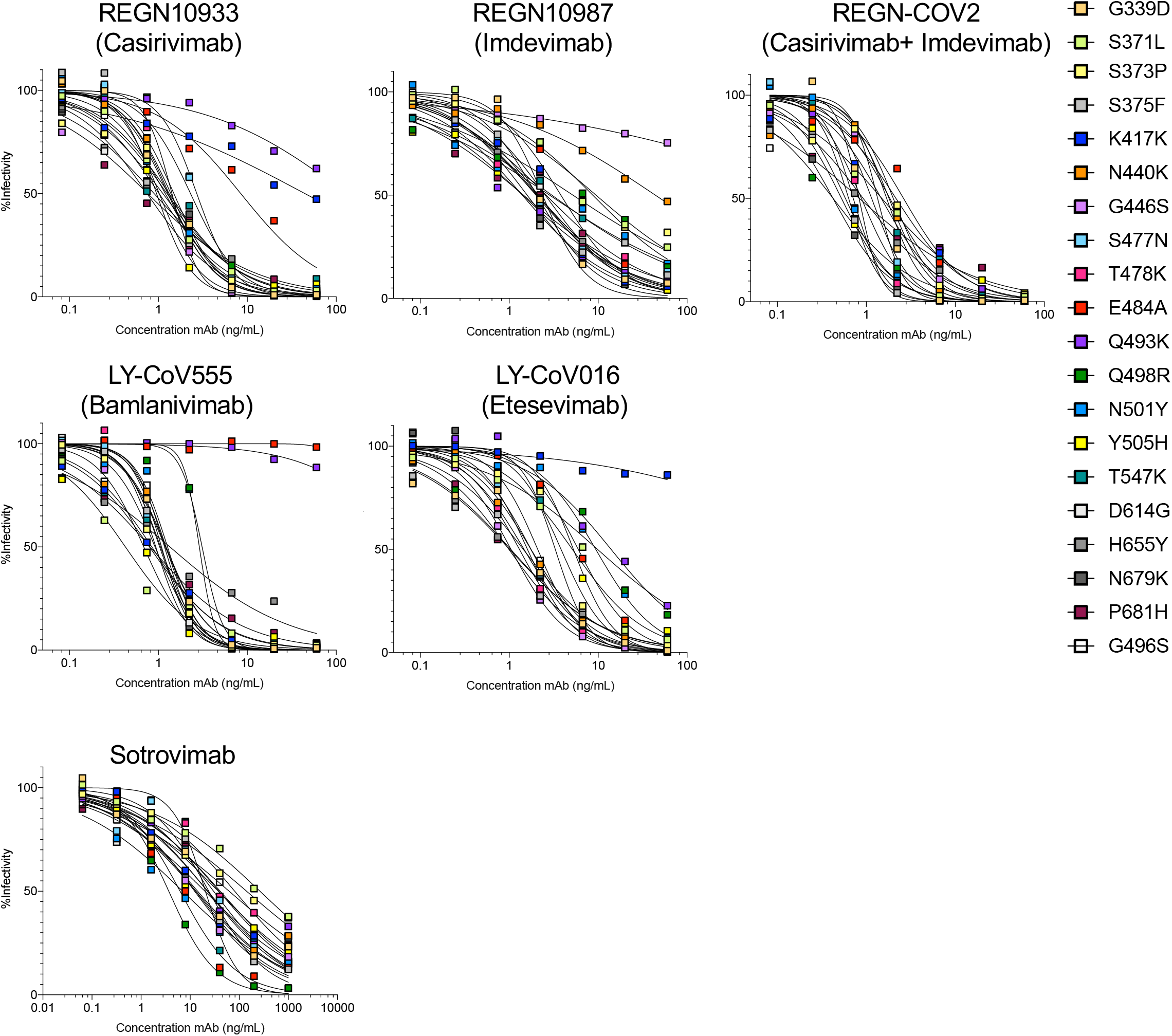
Neutralization of pseudotyped virus with individual mutations by monoclonal antibodies. Neutralization curves were generated for the monoclonal antibodies (Casirivimab, Imdevimab, Bamlanivimab, Etesevimab and Sotrovimab) on viruses pseudotyped by spike proteins with the individual Omicron RBD and cleavage site mutations. All of the spike proteins tested contain the D614G mutation. Neutralization curves were generated for the monoclonal antibodies on viruses pseudotyped by spike proteins with the individual Omicron RBD mutations. The mutations tested include all of the mutations in the RBD and three carboxy-terminal mutations in S1 (H655Y, N679K and P681H).

