## Supplementary material for "Increased resistance of SARS-CoV-2 Omicron Variant to Neutralization by Vaccine-Elicited and Therapeutic Antibodies": Table S1

**Table S1**. IC50 (Figure 1B) of Convalescent, BNT162b2 and mRNA-1273 elicited antibodies against viruses with variant spike proteins. Age, Sex and Comorbidities are shown.

| | **Convalescent** | | | |
|---|---|---|---|---|
| | $IC_{50}$ (serum dilution) | | | |
| donor | D614G | Beta | Delta | Omicron |
| 1 | 311 | 64 | 134 | ND |
| 2 | 239 | 47 | 156 | 16 |
| 3 | 151 | 50 | 40 | ND |
| 4 | 145 | 2 | 101 | 21 |
| 5 | 61 | 36 | 97 | ND |
| 6 | 249 | 130 | 111 | ND |
| 7 | 409 | 119 | 156 | ND |
| 8 | 243 | 43 | 53 | 39 |
| 9 | 217 | 6 | 46 | 5 |
| 10 | 310 | 151 | 122 | ND |
| Mean (SD) | 233 (94) | 65 (49) | 102 (41) | 9 (36) |

| **BNT162b2** | | | | | | | | |
|---|---|---|---|---|---|---|---|---|
| | | | | | $IC_{50}$ (serum dilution) | | | |
| donor | Days post 2nd dose | Age | Sex | Comorbidities | D614G | Beta | Delta | Omicron |
| 1 | 84 | 39 | F | None | 910 | 112 | 211 | 22 |
| 2 | 52 | 23 | F | None | 1573 | 141 | 398 | 11 |
| 3 | 101 | 26 | F | Asthma | 1748 | 463 | 266 | 118 |
| 4 | 109 | 33 | F | None | 1010 | 300 | 642 | 18 |
| 5 | 60 | 35 | F | Hypothyroidism, Psoriasis | 330 | 94 | 421 | 35 |
| 6 | 81 | 42 | F | Asthma | 452 | 65 | 247 | ND |
| 7 | 108 | 26 | F | None | 674 | 195 | 124 | 5 |
| 8 | 107 | 24 | M | None | 548 | 86 | 274 | 11 |
| 9 | 110 | 35 | M | None | 490 | 96 | 338 | ND |
| Mean (SD) | 90 (22) | 31 (7) | | | 859 (476) | 172 (123) | 325 (142) | 25 (36) |

| mRNA-1273 | | | | | | | | |
|---|---|---|---|---|---|---|---|---|
| | | | | | $IC_{50}$ (serum dilution) | | | |
| donor | Days post 2nd dose | Age | Sex | Comorbidities | D614G | Beta | Delta | Omicron |
| 1 | 89 | 26 | M | None | 1530 | 208 | 273 | 94 |
| 2 | 92 | 53 | M | None | 1206 | 59 | 567 | ND |
| 3 | 61 | 67 | M | Prediabetes | 1059 | 110 | 246 | 34 |
| 4 | 93 | 33 | F | None | 757 | 147 | 541 | ND |
| 5 | 44 | 32 | M | None | 716 | 137 | 259 | 93 |
| 6 | 100 | 29 | F | None | 1124 | 489 | 239 | ND |
| 7 | 52 | 33 | F | None | 1074 | 49 | 336 | 42 |
| 8 | 105 | 55 | F | Asthma | 523 | 100 | 123 | 45 |
| Mean (SD) | 80 (24) | 41 | | | 999 (299) | 162 (132) | 323 (145) | 38 (36) |
