## Supplementary material for "Increased resistance of SARS-CoV-2 Omicron Variant to Neutralization by Vaccine-Elicited and Therapeutic Antibodies": Table S2

**Table S2**. IC50 (Figure 1C and D) of BNT162b2 elicited antibodies against viruses with variant spike proteins. Sera were collected from COVID-19 unexperienced and experienced donors. Age, Sex and Comorbidities are shown.

| BNT162b2-COVID-19 unexperienced | | | | | | | | | | | | | | | | | | | |
|---|---|---|---|---|---|---|---|---|---|---|---|---|---|---|---|---|---|---|---|
| | | | | D614G | | | | Beta | | | | Delta | | | | Omicron | | | |
| Donor | Age | Sex | Comorbidities | Pre-vaccination | 1 month post vax-2 | 7-8 month post vax-2 | 1 month post booster | Pre-vaccination | 1 month post vax-2 | 7-8 month post vax-2 | 1 month post booster | Pre-vaccination | 1 month post vax-2 | 7-8 month post vax-2 | 1 month post booster | Pre-vaccination | 1 month post vax-2 | 7-8 month post vax-2 | 1 month post booster |
| 1 | 34 | M | None | ND | 1074 | 165 | 4672 | ND | 227 | 97 | 733 | ND | 187 | 221 | 1060 | ND | 27 | 1 | 300 |
| 2 | 29 | F | None | ND | 564 | 319 | 3561 | ND | 296 | 180 | 1344 | ND | 309 | 250 | 3641 | ND | 98 | 54 | 347 |
| 3 | 37 | F | Allergy | ND | 624 | 283 | 4203 | ND | 410 | 294 | 1359 | ND | 801 | 462 | 4867 | ND | 155 | 63 | 169 |
| 4 | 62 | M | Hypertension, Hyperlipidemia | ND | 368 | 185 | 3361 | ND | 320 | 158 | 2789 | ND | 541 | 325 | 5654 | ND | 35 | 15 | 525 |
| 5 | 39 | M | None | ND | 1413 | 1096 | | ND | 158 | 193 | | ND | 720 | 579 | | ND | 7 | ND | |
| 6 | 37 | F | None | ND | 2538 | 406 | | ND | 499 | 347 | | ND | 1668 | 596 | | ND | 20 | 185 | |
| 7 | 52 | F | Hypertension | ND | 1542 | 852 | 8063 | ND | 285 | 235 | 3986 | ND | 908 | 470 | 3152 | ND | ND | ND | 363 |
| 8 | 34 | M | Hypothyroidism | ND | 534 | 535 | 3141 | ND | 686 | 397 | 1569 | ND | 926 | 230 | 3796 | ND | 57 | 76 | 366 |
| 9 | 43 | M | None | ND | 2480 | 179 | | ND | 404 | 254 | | ND | 746 | 879 | | ND | 30 | 56 | |
| 10 | 38 | F | Asthma, Anemia, Tinea versicolor | ND | 748 | 238 | 7492 | ND | 184 | 124 | 920 | ND | 770 | 631 | 4976 | ND | 56 | 76 | 506 |
| 11 | 38 | M | None | ND | 202 | 1609 | 737 | ND | 780 | 393 | 1904 | ND | 1073 | 585 | 6109 | ND | 83 | 51 | 391 |
| 12 | 52 | F | None | ND | 1368 | 1520 | 8796 | ND | 501 | 404 | 2217 | ND | 1202 | 450 | 4950 | ND | 159 | 106 | 276 |
| Mean (SD) | 41 (9) | | | | 1121 (743) | 616 (505) | 4892 (2517) | | 396 (185) | 256 (105) | 1869 (959) | | 821 (377) | 473 (188) | 4245 (1445) | | 61 (51) | 57 (50) | 360 (104) |

| | | BNT162b2-COVID-19 experienced | | | | | | | | |
| --- | --- | --- | --- | --- | --- | --- | --- | --- | --- | --- |
| | | D614G | | | | Beta | | | | |
| Sex | Comorbidities | Pre-vaccination | 1 month post vax-2 | 7-8 month post vax-2 | 1 month post booster | Pre-vaccination | 1 month post vax-2 | 7-8 month post vax-2 | 1 month post booster | Pre-vaccination |
| M | Asthma | 107 | 3046 | 1321 | 19136 | 19 | 2994 | 2033 | 17987 | 40 |
| F | None | 32 | 2218 | 1885 | 26367 | ND | 2084 | 806 | 20458 | 40 |
| F | Hypertension, Obesity | 41 | 6066 | 4301 | 14407 | 28 | 2132 | 843 | 7861 | 31 |
| M | Cardiovascular disease | 64 | 2137 | 1752 | 7239 | ND | 845 | 1045 | 5420 | ND |
| F | None | 84 | 5849 | 4910 | 14302 | 49 | 1814 | 853 | 10805 | ND |
| F | Diabetes, Herpes simplex | 56 | 6493 | 4721 | 11001 | 25 | 1517 | 1138 | 6501 | 96 |
| M | None | 8 | 3895 | 4003 | 11623 | ND | 2811 | 835 | 5640 | ND |
| | | 56 (31) | 4243 (1734) | 3270 (1435) | 14868 (5790) | 30 (11) | 2028 (683) | 1079 (406) | 10668 (5701) | 52 (26) |
